# Antibiotic-Induced Morphological Changes Enhance Phage Propagation: A Mathematical Model of Plaque Formation in Structured Environments

**DOI:** 10.1101/2024.09.11.612426

**Authors:** Julián Bulssico, Swapnesh Panigrahi, Nicolas Ginet, Mireille Ansaldi

## Abstract

A distinctive manifestation of phage infection in solid media is the appearance of lysis plaques, which correspond to the circular thinning of a bacterial lawn. During plaque formation, successive cycles of phage replication generally take place from a single point of infection and spread radially in a matrix of immobilized bacterial hosts. Many different factors affect plaque size, such as the composition and the reticulation of the propagation matrix, the characteristics of the phage, but also parameters related to the physiology of the bacterial host.

Since combined administration of both antibiotics and phages is a common practice during compassionate treatments, our research focuses on the effects of antibiotics on phage predation, which can be of crucial importance for the therapeutic applications of phages.

It has been previously observed that the presence of antibiotics at sublethal concentrations can affect drastically bacterial physiology, allowing phages to spread more rapidly and resulting in better bacterial eradication. Previous experimental work has focused on the phage characteristics that influence plaque size. However, as plaque formation is strongly influenced by host growth dynamics, a comprehensive model integrating both the host growth and phage infection parameters is required.

In this work, we suggest that plaque enlargement is linked to morphological changes of the host that have an impact on the rate of epidemic propagation and certainly on phage diffusion into the matrix. To support this hypothesis, we characterized the growth parameters of two different phages and bacteria in semi-solid media in the presence of various antibiotics. By combining these data, we have produced a mathematical model that accounts for these observations and explains the increase in plaque size when the host morphology is affected.

**Significance Statement:** This study provides new insights into the phenomenon of Phage-Antibiotic Synergy (PAS) by demonstrating that antibiotic-induced morphological changes in bacterial hosts play a critical role in enhancing phage propagation in structured environments. By linking these morphological alterations—such as cell filamentation and bloating—to increased lysis plaque size, the research underscores the importance of host dynamics in phage therapy. The development of a mathematical model integrating both host growth and phage infection parameters offers a novel framework for understanding and optimizing phage-based treatments in the presence of antibiotics, potentially improving therapeutic outcomes.

## Introduction

Bacteriophages, viruses that parasitize and kill bacteria, are ubiquitous players in every ecosystem on Earth, participating in fundamental biogeochemical processes and driving bacterial evolution through the many complex interactions they entertain with their hosts (1–3). Yet phage research under laboratory settings is fundamental to gain insight on such dynamics as well as for the development of phage therapy to fight against bacterial pathogens.

When culturing bacteria in solid media, one of the most distinctive manifestations of phage infection is the appearance of a lysis plaque, the circular clearing on a bacterial lawn following phage predation. In fact, the term bacteriophage (literally “bacteria eater”) was coined after the observation of this dramatic phenomenon more than a century ago (4). During plaque formation, successive cycles of phage replication take place, usually starting from a single infection spot, and propagating radially on a matrix of immobilized bacterial hosts (5). Many factors are considered to affect plaque size such as the composition of the propagation matrix and the characteristic of a given phage, but also parameters related to bacterial host physiology (6). From a therapeutic point of view, maximizing phage propagation can be an advantage, since it allows the spread within larger infected areas and the predation of larger amounts of bacteria in spatially structured environments such as plant or animal tissues. In this context, it has been documented that the presence of certain antibiotics can boost phage propagation, a phenomenon named Phage-Antibiotic Synergy (PAS) (7, 8). In fact, PAS was known for a long time and in the past two decades its study underwent extensive revival due to its relevance for phage applications. In therapy, combined treatments including both phages and antibiotics are frequent in the case of compassionate care and led to the publication of promising case studies (9–11). Under PAS conditions, bacterial physiology is altered by the presence of antibiotics, allowing for a more efficient propagation of phages. When this propagation takes place in a lawn of bacteria spread on solid medium, synergy can be seen as an increase in the speed of lysis plaque formation, resulting in a larger plaque size. In past publications, synergy was mainly attributed to an increase in phage fecundity, a parameter represented by phage burst-size, which is the number of virions produced per cell (7, 12). However, since plaque formation is strongly influenced by host growth dynamics, a comprehensive model that incorporates host growth and phage infection parameters was still needed. This aspect should not be overlooked in the context of PAS, where the host metabolism and physiology are deeply affected by the presence of a given antibiotic. In this work, we characterized the effects of various antibiotics and found that antibiotic-induced morphological are key to increase the speed of propagation of phage epidemics. In addition to filamentation, which has been correlated with PAS (7, 12–14), we assessed the role of mecillinam-mediated cell bloating in increasing lysis plaque size. Furthermore, by emulating these two altered morphologies via inhibition of key regulators of bacterial shape, we reproduced the increase in lysis plaque size observed during synergy between phages and antibiotics. Our experimental data therefore suggest that the underlying mechanism of PAS is highly dependent on the type of morphological alteration induced by the drugs. We conclude this work with a mathematical model that captures such observations and explains the increase in plaque size observed in the presence of antibiotics. To our knowledge, this is the first experimental work to characterize epidemic dynamics in the context of PAS, taking into account morphological changes associated to antibiotics.

## Results

### Plaque size increases in the presence of sublethal doses of antibiotics that modify cell shape

To approach PAS in a systematic way, we tested if several antibiotics belonging to different families could increase the radii of lysis plaques of phages T5 and T7 infecting *E. coli* MG1655. The antibiotics selected in this study cause either bacterial filamentation by impairing division (ciprofloxacin and ceftazidime), cell bloating (mecillinam), or no effect on the cell morphology (kanamycin and chloramphenicol). In order to measure the influence of cell shape modification on the size of plaques generated by phages, we measured the radii of plaques formed by phages T5 and T7 in the presence of sublethal doses of the antibiotics mentioned above.

Figures 1a and 1b show the effect of sublethal antibiotic concentrations on the aspect of lysis plaques of phages T5 and T7, respectively. These plaques were obtained through a standard agar overlay assay in the presence or absence of drugs. For each antibiotic, the highest concentration tested corresponds to its maximum sublethal dose (see Material & Methods) which does not lead to a significant reduction in the density or homogeneity of the bacterial lawn in the top agar. For phages T5 and T7 the only antibiotics that induced a significant increase in plaque size were ciprofloxacin, ceftazidime, and mecillinam, whereas neither chloramphenicol nor kanamycin showed such effect.

**Figure 1.**
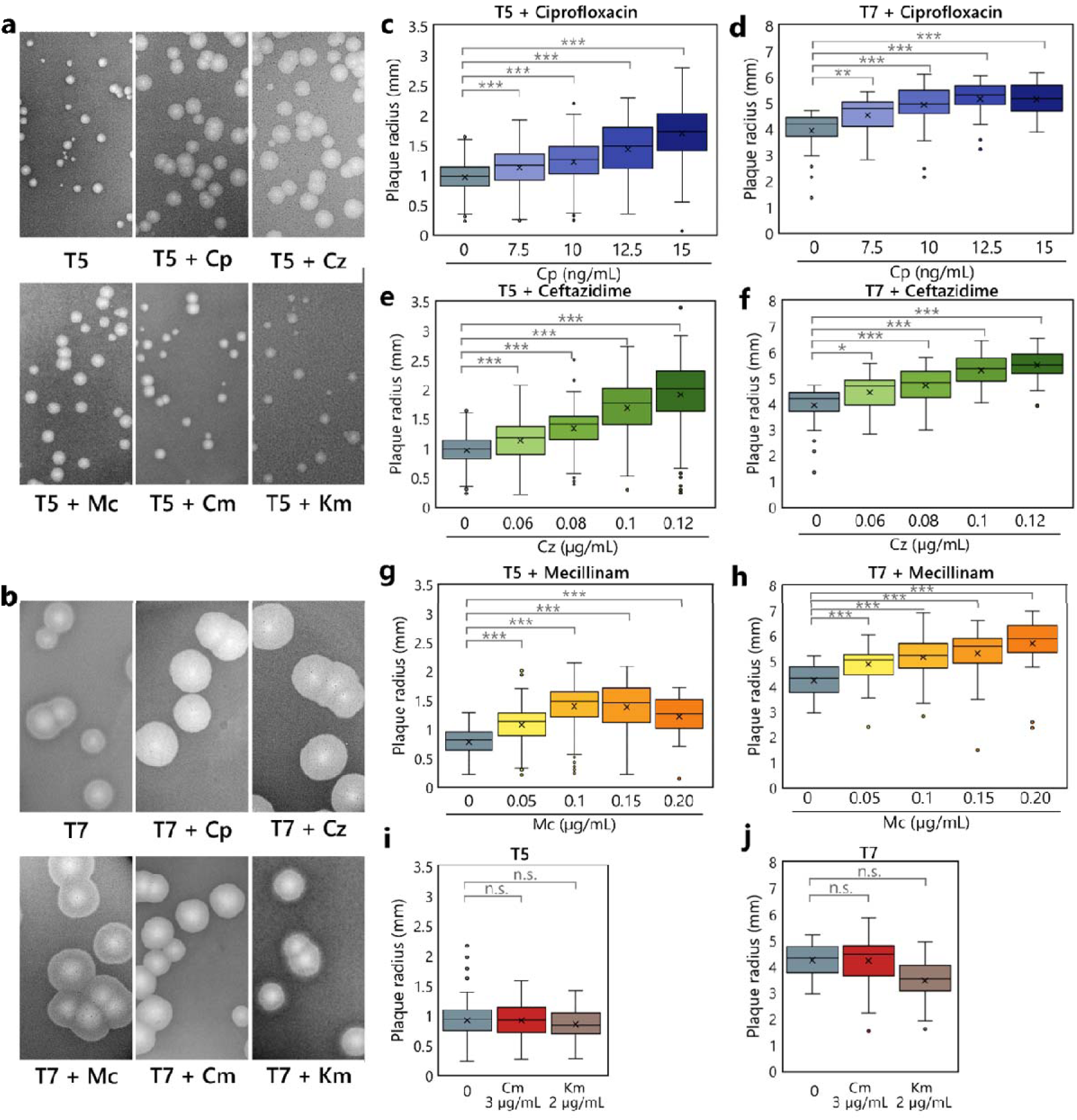
Lysis plaques produced by phages T5 and T7 on *E. coli* MG1655 are enlarged in the presence of sublethal doses of antibiotics. (**a, b**) Aspect of the lysis plaques of phages T5 and T7 at the maximum sub-inhibitory concentration of the tested antibiotics. (**c, d, e, f, g, h, I, j**) Boxplots of lysis plaque radius measurements at increasing concentrations of: ciprofloxacin (Cp), ceftazidime (Cz), mecillinam (Mc), chloramphenicol (Cm) and kanamycin (Km) for phages T5 and T7. The gradient of concentration used was right below the inhibitory concentration. N = for each condition, between 18 and 414 plaques were measured. Whiskers represent the range of the data within 1.5 times the interquartile range from the lower quartile and upper quartile, mean values are shown as crosses. Three asterisks represent a p-value of less than 0.001 for an upper-tailed t-test comparing between treated and untreated conditions, two asterisks represent a p-value of less than 0.01, and one asterisk represent a p-value of less than 0.05; whereas “n.s.” represents non-significative results, this is, p-values of more than 0.05 in a same test.

In order to measure this phenomenon and to determine if the synergistic behavior between phages and antibiotics was dose-dependent, we carried out similar agar overlay assays across a gradient of increasing antibiotic concentrations, measuring the radii of the resulting plaques. The mean radii of lysis plaques produced by phages T5 and T7 increased in the presence of the two filamentation-inducing antibiotics, namely ciprofloxacin and ceftazidime in a dose-dependent manner, compared to the untreated condition (Fig. 1). Phage T5, which has a mean plaque radius of 0.88 ± 0.26 mm after 24 hours of incubation in the absence of antibiotics, reached a mean radius of 1.70 ± 0.44 mm (93% increase) and 1.91 ± 0.57 mm (117% increase) in the presence of 15 ng/ml ciprofloxacin and 120 ng/ml ceftazidime, respectively. Similarly, T7 lysis plaques with a mean radius of 4.12 ± 0.69 mm after 24 hours of incubation, showed a maximum radius of 5.13 ± 0.69 mm (25% increase) and 5.49 ± 0.62 mm (33% increase) with 15 ng/ml ciprofloxacin and 120 ng/ml ceftazidime, respectively. Furthermore, a gradual increase in the concentration of mecillinam, which induces cell bloating, also correlated with an increase in the average plaque radius of both phages (Fig. 1g, 1h), with a maximum radius of 1.38 ± 0.41 mm (57% increase) and 5.69 ± 0.98 mm (38% increase), for phages T5 and T7 respectively. Hence, we conclude to a dose-dependent synergistic effect of both ciprofloxacin and ceftazidime with two phage species belonging to two taxonomically distant phage families (*Demerecviridae* for T5 and *Autographiviridae* for T7). In contrast, no significant increase in plaque size was observed at the maximum sublethal concentration of chloramphenicol and kanamycin for both phages (Fig. 1i, 1j), evidencing the absence of antibiotic effect of chloramphenicol on phage propagation in both cases. In contrast, kanamycin added at the maximum sublethal concentration highlights the specificity of phage / antibiotic combination with the absence of effect on plaque size for T5 and a reduction for phage T7 suggesting an antagonistic interaction. Such antagonistic interactions between phages and some antibiotics have already been documented (15, 16). Altogether, these results highlight a strong correlation between the increase in propagation speed of phages T5 and T7, and the presence of certain types of antibiotics, notably ciprofloxacin, a DNA gyrase inhibitor and known inducer of cell filamentation through the activation of the SOS response, cefalexin and mecillinam, which both belong to the β-lactams family.

### Two morphological changes, filamentation and cell bloating, correlates with an increase in plaque size

To evaluate the extent of morphological changes in *E. coli* cells triggered by the presence of the synergistic drugs in our experimental conditions, we measured morphological changes of individual bacteria in cultures treated with increasing sublethal antibiotic concentrations were measured. Figure 2a showcases phase contrast microscopy images of bacterial microcolonies exposed to the previously described antibiotics. Morphological changes such as cell filamentation (ciprofloxacin and ceftazidime) or cell bloating (mecillinam) were observed and increased in a dose-dependent manner. As expected, control antibiotics (chloramphenicol and kanamycin) did not provoke any morphological modifications. To quantify these observations, large populations of bacteria were imaged and measured under the same drug concentrations. Figure 2b scatterplot summarizes such effects through two morphological descriptors: mean cell width and mean cell length. As previously observed, the antibiotic ceftazidime increased cell length in a dose-dependent manner, without significantly modifying cell width. The increment on average length observed at the highest concentration of the drug was of 550% compared to the untreated condition. Ciprofloxacin also increased cell length in a dose-dependent manner with a maximum increase of 300%. Additionally, ciprofloxacin moderately impacted width, with a maximum increase of 10%. On the other hand, mecillinam caused an increase in cell width progressively with increasing doses. At the highest sublethal dose (200 ng/mL), we observed a maximum increase in mean cell width of 36%, without affecting significantly cell length (Fig. 2b). Finally, neither chloramphenicol nor kanamycin significantly affected mean cell length or width, even at the highest sublethal concentrations (3 and 2 µg/mL, respectively). Taken together, our results showed that the degree of morphological change correlated with the strength of synergy measured by plaque size radii.

**Figure 2.**
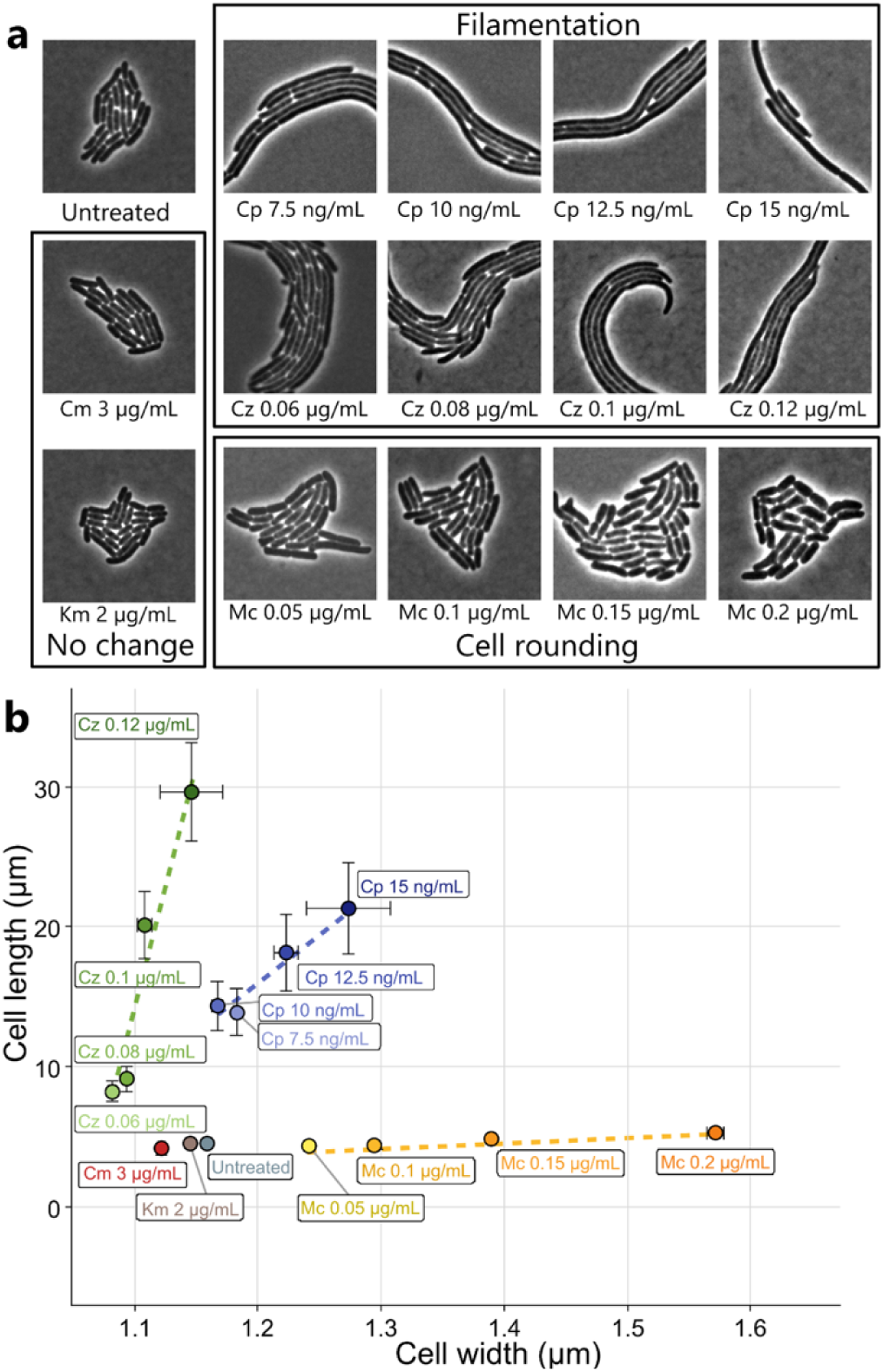
Morphological effects of sublethal antibiotics in *E. coli* MG1655. **(b)** Phase-contrast microscopy of *E. coli* microcolonies trapped in a bidimensional LB-agarose 1% pad, in the presence of the same antibiotic concentrations tested in Fig 1. **(a)** Scatter plot showing the mean bacterial cell width and length of *E. coli* populations treated with increasing concentrations of ciprofloxacin (Cp), ceftazidime (Cz), mecillinam (Mc), chloramphenicol (Cm) and kanamycin (Km). N = for each condition, between 171 and 2684 individual bacteria were measured. Whiskers represent the standard error of the mean.

### Changes in morphology alone are sufficient to increase the size of T5 and T7 lysis plaques on E. coli

Our results suggest that two completely different morphological changes, filamentation and cell rounding, correlate with an increase in the mean radius of lysis plaques. To demonstrate whether filamentation or cell bloating alone were sufficient to induce an enlargement of phage lysis plaques, we aimed at emulating these morphological perturbations in *E. coli* without using the antibiotics mentioned earlier. To achieve this, we resorted to a system based on the “dead” Cas9 (dCas9) enzyme, a variant of the well-known Cas9 from *S. pyogenes* inactivated in its endonuclease activity (17). This variant is still able to use a sgRNA to interrogate and bind to the target DNA. However, due to its abolished catalytic activity, the dCas9-sgRNA complex remains bound to the target dsDNA molecule without cleaving it (18). If the sgRNA is designed to target a particular promoter or the coding sequence of the downstream ORF, the binding of the dCas9 will physically block the gene’s transcription by preventing RNA polymerase binding or elongation. The dCas9 encoding gene is carried by the *E. coli* strain LC-E75, which is an *E. coli* MG1655 derivative encoding the *dcas9* gene under the control of a promoter inducible by anhydrotetracycline (ATc), whereas the sgRNA is constitutively produced from the plasmid psgRNA. The specificity of the targeted gene was conferred by the 20 complementary nucleotides cloned within the sgRNA on the plasmid (see Materials and Methods). We inferred that by impairing the expression of key genes involved in cell morphology we could mimic cell elongation or cell rounding triggered by the above-mentioned antibiotics. FtsZ is a major component of the division machinery in many bacterial species and its depletion blocks cell septation and produces long filaments (19), emulating ceftazidime- and ciprofloxacin-induced morphological changes. Likewise, we expected that impairing *mreB* gene expression, that encodes a key component of the bacterial elongasome and leads to round and bloated cells (20), would mimic mecillinam-induced morphological changes. Figure 3a left column shows cell morphology after two hours induction by ATc in the case of a non-targeting sgRNA (upper panel), a *ftsZ*-targeting sgRNA (center panel) and a *mreB*-targeting sgRNA (lower panel). As expected, filamentation and cell bloating were induced when *ftsZ* or *mreB* expression was impaired, respectively. The negative control showed that the expression of *dCas9* alone did not modify cell morphology. These results demonstrated that we could selectively mimic antibiotic-induced cell morphology changes. We thus proceeded to the measurement of lysis plaques with both T5 and T7 as previously described (Fig. 3a, middle and right columns, respectively). With both T5 and T7 we observed that the two dCas9/sgRNA mediated morphological changes correlated with a strong increase in the size of lysis plaques compared with the condition with non-targeting sgRNA, reminiscent of increase observed with ceftazidime, ciprofloxacin and mecillinam. These results thus support the hypothesis that either filamentation or bloating by themselves allowed increased phage predation.

**Fig 3.**
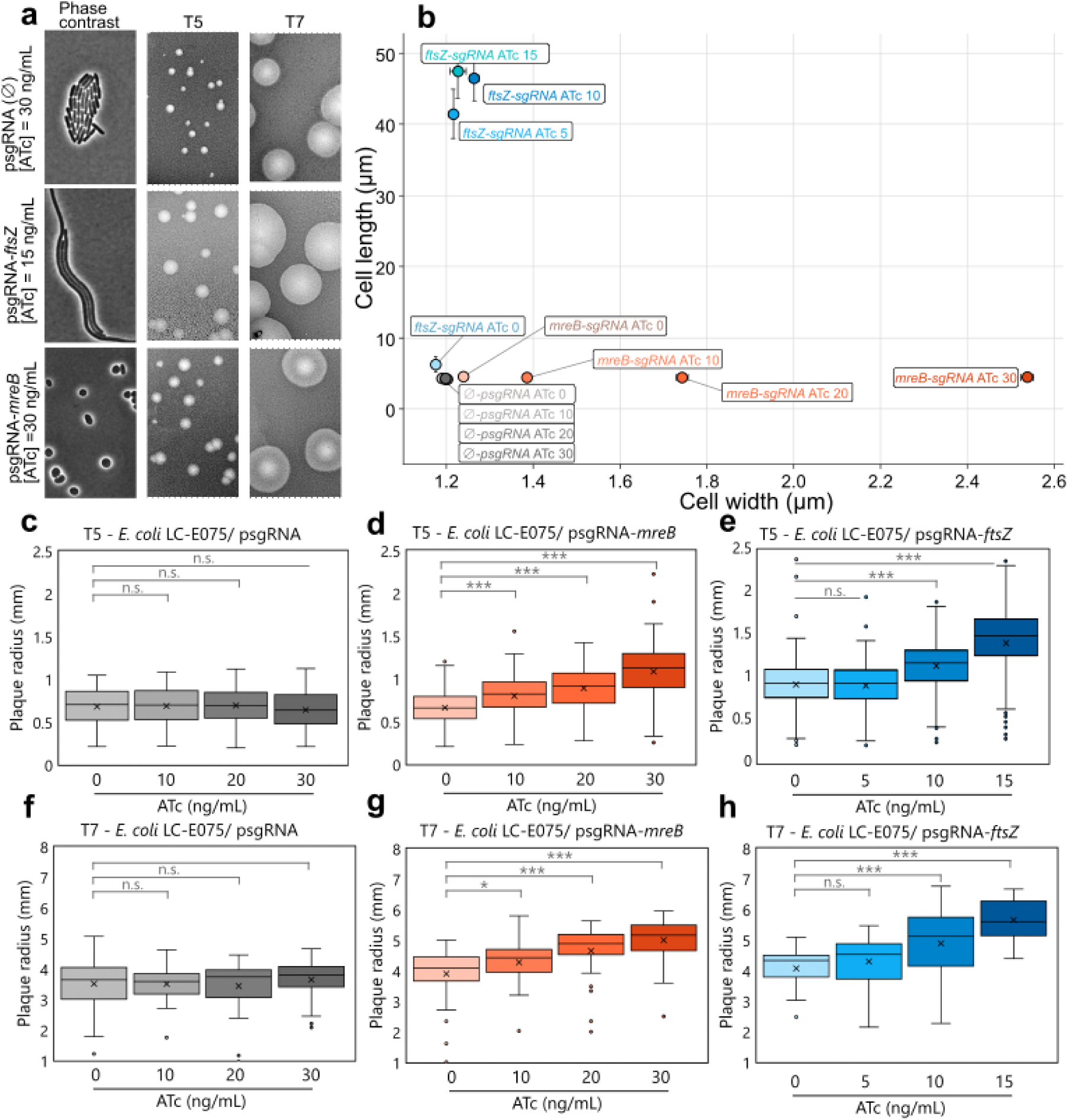
dCas9/sgRNA-driven morphological changes in bacteria produce larger lysis plaques. **(a)** Effect of dCas9/sgRNA repression of key regulators of bacterial morphology and its impact on plaque size of phages T5 and T7. First column: phase-contrast microscopy of *E. coli* LC-E75 microcolonies under an agarose pad. Each image showcases a microcolony of the LC-E75 strain carrying a sgRNA that either does not target any *E. coli* locus (∅), targets *ftsZ* open reading frame or target *mreB* open reading frame, in the presence of ATc at maximum sublethal dose. Second and third column, images of lysis plaques of phages T5 and T7 on LCE75 carrying the same sgRNA and under the same induction concentrations. **(b)** Scatter plot showing the mean bacterial width and length of *E. coli* LC-E75 in the presence of dCas9 and the sgRNAs mentioned above, at increasing inducer (ATc) concentration. N = For each condition, between 36 and 2423 individual bacteria were measured. Whiskers represent the standard error of the mean. **(c, d, e)** Measurements of phage T5 lysis plaque radii on *E. coli* LC-E75 with either a non-targeting sgRNA, sgRNA-*mreB*, or sgRNA-*fstZ*. **(f, g, h)** Measurements of phage T7 on *E. coli* LC-E75 with either a non-targeting sgRNA, sgRNA-*mreB*, or sgRNA-*fstZ*. N = For each condition, between 21 and 352 plaques were measured. Whiskers represent the range of the data within 1.5 times the interquartile range from the lower quartile and upper quartile, mean values are shown as crosses. Three asterisks represent a p-value of less than 0.001 for a upper-tailed t-test comparing between treated and untreated conditions, one asterisk represent a p-value of less than 0.05; whereas “n.s.” represents p-values higher than 0.05 in a same test.

Since we previously observed that cell morphology as well as lysis plaques size vary with increasing concentrations of ceftazidime (cell elongation) and mecillinam (cell bloating) (Fig. 2b), we quantified the effect of *ftsZ* and *mreB* gene repression using the same morphological descriptor (*i.e.*, cell width, cell length and lysis plaque radius) 24 hours post-infection in the presence of increasing concentrations of inducer. Figure 3b shows the average bacterial width and length measured in strains expressing dCas9 with a sgRNA targeting either *ftsZ*, *mreB*, or a control sgRNA that did not target any *E. coli* gene. In the presence of a spacer targeting *mreB* (cell bloating), we observed an ATc dose-dependent increase in cell width reaching a maximum of 2.54 ± 0.44 µm at 30 ng/ml ATc compared to 1.20 ± 0.07 µm in the strain carrying the empty vector at the same induction level (112% increase). On the contrary, the increase in average cell length observed in the presence of an *ftsZ*-targeting spacer (cell filamentation) almost reached its maximum extent with as few as 5 ng/mL of ATc. The maximum elongation was observed at 15 ng/mL of ATc with a mean length of 47.47 ± 26.53 µm, compared to 4.35 ± 1.32 µm in the control experiment with the non-targeting sgRNA (996% increase).

dCas9-induced morphological changes properly imitate the effects of the synergistic antibiotics we tested previously. Firstly, the increase in length obtained with dCas9-mediated repression of *ftsZ* is almost twice as big as in the presence of ceftazidime or ciprofloxacin. Secondly, the increase in bacterial width is a consequence of *mreB* inhibition is three times the one observed with mecillinam. It is likely that the presence of antibiotic disturbs other key cellular processes leading to cell death before achieving the extreme changes in cell morphology observed upon inhibition of *ftsZ* or *mreB* genes.

We thus successfully adapted a dCas9-based tool to reproduce two morphological effects, filamentation and bloating, observed upon sublethal antibiotic treatments. We then wondered if such morphological changes, obtained in the absence of antibiotics, could account for enlarged lysis plaques previously observed in the presence of antibiotics (Fig. 1). To evaluate this, we carried out the standard agar overlay assay as described before with increasing concentrations of the inducer to evaluate the size of the lysis plaques during infection by phages T5 and T7 (Fig 3c-3h). When a non-targeting psgRNA was used, we did not observe any significant increase of the lysis plaque radii either with T5 or T7, whatever the inducer concentrations (Fig 3c, 3f). In the presence of *mreB*-targeting or *ftsZ*-targeting psgRNA, we observed an ATc dose-dependent increase in mean lysis plaque radii for T5 (Figure 3d and 3e) and T7 (Fig 3g, 3h). When cell bloating is induced by repressing *mreB* expression, lysis plaque radii increase by 63% and 28% compared to the control experiment for T5 and T7, respectively (Fig 3d, 3g). When cell filamentation is induced by repressing *ftsZ* expression, lysis plaque radii increase by 53% and 38% compared to the control experiment for T5 and T7, respectively (Fig. 3e, 3h). These results suggest that antibiotic-mediated synergies reported in Figure 1 are chiefly driven by bacterial morphology changes (filamentation for ceftazidime and ciprofloxacin, bloating for mecillinam) and not by other effects of the antibiotic on the host. In other words, our findings suggest that cell bloating or filamentation, induced either by antibiotics of by the use of the dCas9/sgRNA system, would be the main factor responsible for the increase in both T5 and T7 lysis plaques size.

### Filamentation increases phage burst size through different mechanisms

In the previous sections, we used a dCas9/sgRNA system to block the transcription of genes involved in the maintenance of *E. coli* shape in order to demonstrate that both bacterial filamentation and bacterial bloating were sufficient to increase the plaque size of phages T5 and T7. After having established a link between morphological changes and increased plaque size, we questioned the underlying mechanisms of plaque enlargement driven by two contrasting morphological effects. To test this, we should determine whether filamentation and bloating impact phage predation in a similar way or contribute to plaque enlargement through distinct mechanisms.

In order to assess the impact of a given morphological change in the replicative cycle of phages T5 and T7, we carried out one-step growth curve experiments in the presence of dCas9-mediated morphological changes achieved through inhibition of *ftsZ* or *mreB* gene transcription. Figure 4 showcases replication of phages T5 and T7 at 37°C in liquid LB medium. For phage T5 (Fig. 4a) a latent period of 50 minutes and a burst size of 110 PFU/bacterium were observed in the presence of a non-targeting sgRNA, in agreement with the literature (21). When filamentation was induced through the inhibition of *ftsZ* transcription, a delay of about 15 min in the latent period was observed as well as an increase of 30% in burst size. The cause of the increase in progeny size might be attributed to a delay in the period of intracellular phage assembly, which allowed a larger pool of phages to be produced and maturated per cycle. In contrast, cell bloating achieved through *mreB* inhibition neither modified the latent period nor the burst size of phage T5. This suggest that plaque enlargement under bloating is not a consequence of increased phage fecundity.

**Figure 4.**
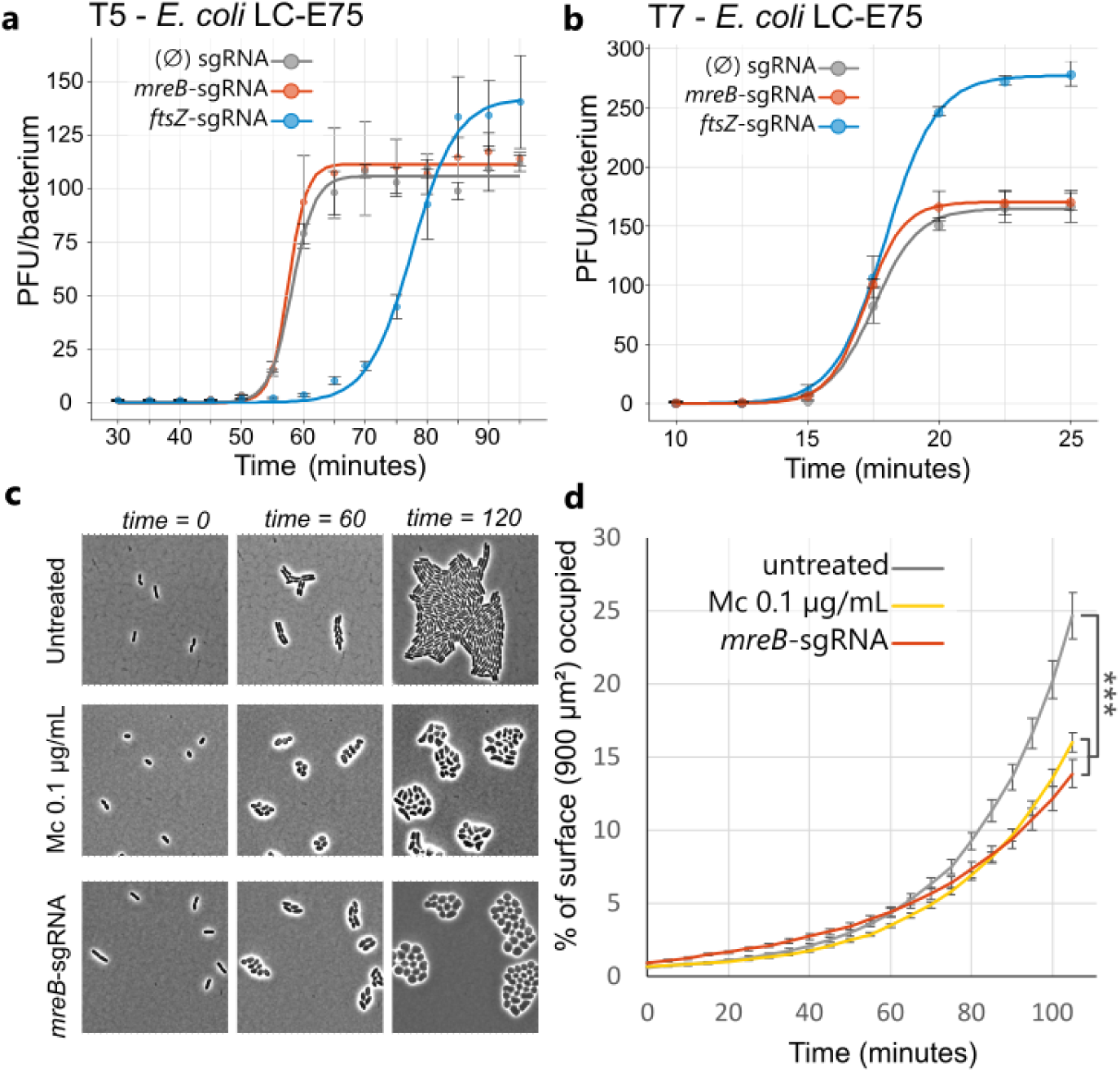
Effects of dCas9-mediated morphological changes on key parameters influencing epidemic spread. **(a,b)** One step growth curve of phage T5, and phage T7 in *E. coli* LC-E75 displaying different morphological changes. Time = 0 represents the time of phage-bacteria mixing. Induction of the morphological change by dCas9 transcription inhibition was carried out for 2 hours prior to the beginning of the infection. Morphological changes at the time of the phage addition were confirmed by phase-contrast microscopy. Concentrations of the inducer were the maximum sublethal concentrations tolerated by the strain. N = 3 curves per condition. Error bars represent the standard error of the mean. **(c)** Phase contrast microscopy images of timelapse growth of bacteria in the presence of mecillinam in *E. coli* MG1655 or after repression of the *mreB* by the dCas9/sgRNA in strain LCE-75. Cells were cultured in liquid medium and pre-treated with the antibiotic or the inducer one-hour prior sampling and microscopy imaging. N = between 30 to 43 microcolonies were followed and measured per condition. **(d)** Percentage of a given surface area occupied by microcolonies of *E. coli* under different conditions over 105 minutes of growth at 37°C. Error bars represent the standard error of the mean. Three asterisks represent a p-value of less than 0.001 for an upper-tailed t-test comparing values at time = 105 minutes.

We also compared replication of phage T7 under the same conditions (Fig. 4b). In the absence of morphological alteration, phage T7 displayed a latent period of 15 minutes and a burst size of approximately 160 PFU/bacterium, which corresponds to expected burst size (21, 22). Upon *ftsZ*-mediated filamentation, burst size increased by 70% compared to the control with a non-targeting spacer. However, unlike phage T5, filamentation did not impact T7’s latent period. The production of a larger phage progeny within a similar latent period could indicate that filamentous bacterial cytoplasm may allow for larger phage factories to be formed prior to lysis. Finally, as observed for phage T5, bacterial bloating caused by *mreB* inhibition neither affected significantly the latent period nor the burst size of phage T7, suggesting a different mechanism accounting for enhanced phage predation.

### Cell bloating improves phage diffusion by reducing phage-bacteria interactions

Since the replicative cycles of phages T5 and T7 were not modified when bacteria suffered cell bloating, we hypothesized that the increase in plaque size was rather due to changes in the dynamics of phage-bacteria interactions during propagation. These changes could improve phage diffusion in the bacterial lawn matrix. To test this hypothesis, we measured the packing of *E. coli* microcolonies either in the presence of mecillinam or upon *mreB* expression inhibition (Fig. 4c, 4d). As observed in Figure 4c, whatever the cause of cell bloating, microcolonies tend to adopt round and compact arrangements over time compared to the untreated condition. To quantify this, we measured the percentage of bidimensional space occupied by each microcolony over time (Fig. 4d): bloated bacteria tend to occupy around 38% less space compared to microcolonies composed of regular-shaped cells after 100 minutes of incubation. The boost of phage propagation can then be explained as follows: a reduction of the fraction of space occupied by the host is predicted to increase the diffusion of viral particles (6). Consequently, in the presence of compact arrangements of bacteria allowed by cell bloating, phages can diffuse and penetrate further in the bacterial matrix before adsorbing to a non-infected host, increasing the overall speed of phage propagation and then the lysis plaque radius.

### Recording phage T7 propagation

In order to study the propagation of phage epidemics we followed the dynamics of phage propagation by recording lysis plaque expansion on plates with time laps imaging. In the case of T7 this can be conveniently carried out over relatively long periods of time due to the phage ability to infect bacterial cultures that have entered into stationary phase (23). We thus monitored T7 lysis plaque formation in the presence of increasing concentrations of mecillinam (inducing cell bloating) and ciprofloxacin (inducing cell filamentation). Figure S1 shows the resulting kinetics of plaque enlargement.

In all tested conditions, these kinetics were biphasic. The first phase lasted about 14 hours and the propagation rate - measured as the slope of plaque formation kinetics - was maximum and fairly constant for all antibiotic concentration tested. This initial phase corresponds to the bacterial lawn exponential growth phase. The shift to the second kinetic regimen follows the entry of the bacterial lawn into the stationary phase. During this second phase the propagation rate decreased compared to the initial phase but in an antibiotic dose-dependent manner for the two tested antibiotics. At variance with the initial phase where propagation rates are globally unaffected by the antibiotic concentrations, we observed during this second phase that rates increased with the antibiotic concentrations. These results are in accordance with increases in plaque size measured at a fixed time post infection we previously observed (Fig. 1). Altogether, these kinetics profiles suggest that the increase in plaque size was mostly due to an accelerated T7 propagation taking place in the mature bacterial lawn stimulated by the addition of mecillinam and ciprofloxacin.

### A mathematical model to study plaque enlargement

#### Theoretical framework

We measured the rate of T7 plaque formation in the presence of mecillinam and ciprofloxacin, two antibiotics that accelerate phage propagation at synergistic antibiotic concentrations, and studied the physiological changes induced in phage and bacteria. We thus came up with a mathematical model that aims to describe the changes in T7 propagation rate as a function of the changes in bacterial and phage physiology. Equations mentioned underneath are detailed in the Supplementary file, and given the assumptions we made, we provide a theoretical description of the square of terminal velocity of lysis plaque expansion speed in each experimental case *c_T_* (T for antibiotic-treated cells) relative to the speed of untreated host *c_u_* (u for untreated cells):

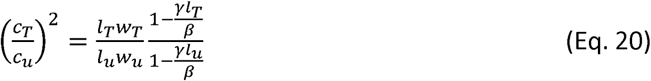

In our theoretical framework, this dimensionless parameter captures the impact of cell morphological changes on phage lysis plaque propagation we previously evidenced (Fig. 1 and Fig. 3).

When cell length *I* remains constant as it is the case with mecillinam (Figure 2) or when *mreB* gene expression is repressed (Fig. 3), Eq. 20 can be simplified as:

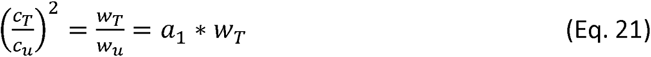

Conversely, when cell width *w* remains constant as it is the case with ciprofloxacin (Fig. 2) or when *ftsZ* gene expression is repressed (Fig. 3), Eq. 18 can be simplified as:

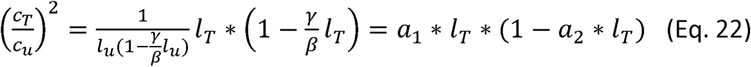

In both cases, the dimensionless parameter describing the dynamics of phage lysis plaque expansion at steady-state is a function of a single cell morphological parameter, either width *w* or length *I*.

#### Experimental data

We recorded lysis plaque radius (*r-r*_0_) enlargement kinetics on solid media for 28 h for untreated cells and cells treated with increasing concentrations of mecillinam (0 – 70 ng/mL) inducing cell bloating (Fig. S1a) or ciprofloxacin (0 – 8 ng/mL) inducing cell filamentation (Fig. S1b). As mentioned earlier, these kinetics are biphasic with a rapid initial plaque expansion followed by a slower after approx. 16h. We defined for each kinetics the terminal velocity *c* (*c_T_* for treated cells and *c_u_* for untreated cells) as the slope of slow phase derived from the experimental kinetics. We then plotted the squared relative terminal velocity 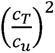 against cell width w for mecillinam (Fig. 5a) and against cell length I for ciprofloxacin (Fig. 5b).

**Figure 5:**
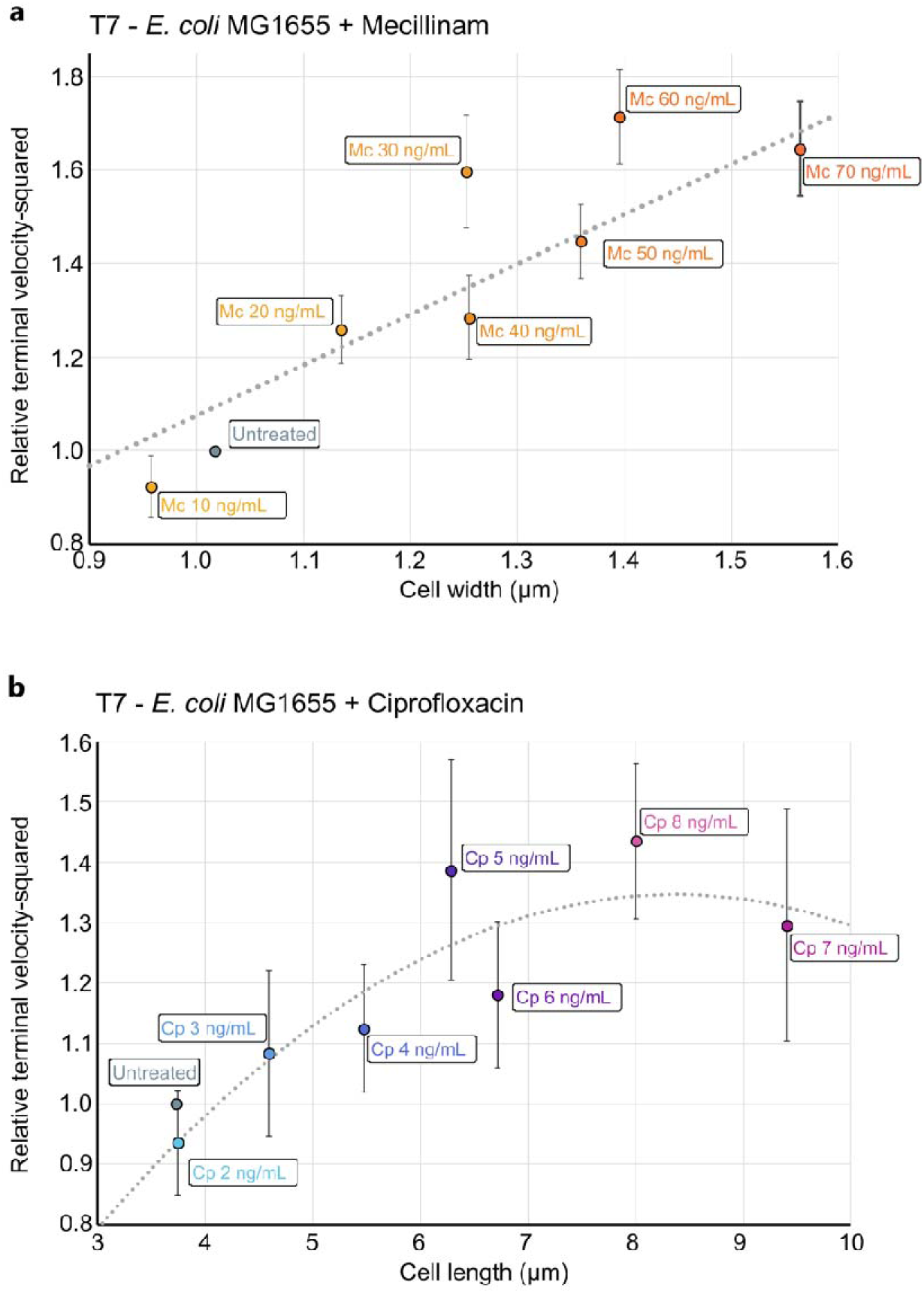
Square relative speeds of different concentrations of mecillinam and ciprofloxacin is plotted, respectively, against width and length of cells. The theoretical model leads to the 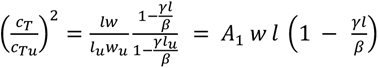. In case of cells treated with mecillinam the length of cells did not change while the width changed leading to a linear relation with width as shown in [a], where the dots represent the experimentally determined speed as a function of cell width. Similarly, for cells treated with ciprofloxacin the lengths change keeping the width same. The experimentally determined relative square speed is then plotted (dots) as a function of length and fit is shown.

According to Eq. 21, in the presence of mecillinam we expect that 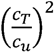 increases proportionally with cell width w. Experimental data in Figure 5a can indeed be satisfactorily fitted with the linear function *y* = *a*_1_.*x* with *a*_1_ = 1.08 ± 0.58 μ*m*^-1^.

According to Eq. 22, in the presence of ciprofloxacin we expect that 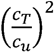 follows a second-degree polynomial equation, increases with cell length *l* and reaching a maximum at *l*= *β*/2*γ* then decreasing while cells keep elongating. Experimental data in Figure 5a does exhibit such a trend and were fitted with the second-degree polynomial function *y* = *a*_1_\**x*\*(1-*a*_2_\**x*) with 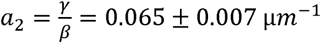 and *a*_1_ = 0.349± 0.018 μ*m*^-1^. The fitting curve reaches a maximum at *l* = 7.71 μ*m*.

In both mecillinam and ciprofloxacin cases, our theoretical model is in accordance with the experimental data. For a given set of experimental conditions, PAS efficiency measured through phage lysis plaque expansion kinetics is governed by cell morphological changes (length or width) induced by the antibiotic treatment administrated in conjunction with the phage treatment. We can predict that PAS efficiency keeps increasing while cell width increases (*e.g.*, mecillinam, Fig. 5a) but reaches an optimum while cell length increases (*e.g.*, ciprofloxacin Fig. 5b).

This model yields several predictions on PAS efficiency when other key parameters related to phage or cellular cycles vary (Fig. S2). Thus, for a given antibiotic concentration, lysis plaque expansion speed and radius are expected to increase when phage adsorption rate a decreases (Fig. S2c). Varying phage inactivation rate κ (assumed to be very small in our model and for phage T7) has very little effect on lysis plaque expansion dynamics and radius. The terminal velocity is not affected and we just expect a small decrease in lysis plaque radius when κ increases (Fig. S2d).

External parameters affecting biomass growth rate (*e.g.*, nutrient availability) have a severe impact on lysis plaque radius with increased lysis plaque size when *µ* decreases in growth limitation conditions. Saturation is reached faster when *µ* increases (Fig. S2e). Finally, biomass capacity increases, both radius and expansion speed are decreased.

We thus provide an explainable model that fits with experimental data related to cell morphological parameters. From this model we could predict the effect of parameters related to phage cycle and external parameters affecting cell growth on lysis plaque expansion speed and radius.

## Discussion

The objective of the present study was to characterize the synergy taking place between two taxonomically distant bacteriophage species (T5 and T7) and several antibiotics with contrasted effects on bacterial morphology. The rationale was to determine whether synergy can take place through a common mechanism for different antibiotic classes, or on the contrary through distinct effects in the phage-bacteria system. To achieve this, we used different classes of antibiotics, namely β-lactam (ceftazidime, a cephalosporin and mecillinam, an aminopenicillin), fluoroquinolone (ciprofloxacin), aminoglycoside (kanamycin) and chloramphenicol. We measured phage-antibiotic synergy (PAS) as the increase of the size of T5 and T7 lysis plaques on lawns of *E. coli* MG1655 undergoing antibiotic stress. The range of antibiotic concentrations we used was right below the threshold of growth inhibition of the bacterial lawn. Under these conditions, we first observed that the tested antibiotics could be classified into two families regarding PAS (Fig. 1). The first one (ceftazidime, ciprofloxacin and mecillinam) regroups antibiotics eliciting a dose-dependent enlargement of phage lysis plaques compared to the untreated condition. The second one (kanamycin and chloramphenicol) regroups antibiotics that do not modify the size of the phage lysis plaques. Results were consistent for both bacteriophages T5 and T7 although they belong to taxonomically distant families (*Demerecviridae* for T5 and *Autographiviridae* for T7). We thus successfully set up a standardize method to screen and quantify the PAS effect for various combinations of phage / antibiotic.

Since we previously evidenced that cell filamentation is a key driver of successful phage predation in the PAS context we examined the consequences of morphological alteration of bacterial cells on phage predation in a broader context (13). Indeed, the antibiotics we tested can now be classified into three families (Fig. 2). The first one comprising ciprofloxacin and ceftazidime induces *E. coli* cell filamentation. The second one comprising mecillinam induces cell rounding. Both phenomena take place in a dose-dependent manner and in the exact same range of concentrations where phage plaque enlargement occurs (Fig. 3). The third family including kanamycin and chloramphenicol does not elicit any cell morphological change. One has to note that the three synergistic antibiotics (ceftazidime, ciprofloxacin and mecillinam) belong to two different classes (β-lactam and fluoroquinolone) and that two molecules belonging to the same class (β-lactam) triggers opposite morphological changes (cell filamentation for ceftazidime and cell bloating for mecillinam). Since in the absence of antibiotic-induced morphological changes no synergistic effect is observed, we hypothesize a direct correlation between morphological alteration and accelerated phage propagation rather than with the direct molecular mechanisms specific to each antibiotic.

In order to prove that filamentation and cell rounding were necessary and sufficient to enhance phage propagation, we used a dead-Cas9 system to trigger such altered morphologies. By downregulating the expression of key genes responsible for the maintenance of *E. coli* cell shape (*ftsZ* for cell filamentation and *mreB* for cell rounding), we successfully emulated cell filamentation and cell rounding without the requirement of antibiotics. Strikingly, the effects of cell morphology alteration on phage predation were completely equivalent to those observed in the presence of the antibiotics and proportional to the extent of the morphological changes. Through this, we confirmed that synergy happen just by modifying bacterial shape into filaments or spheres.

The link between bacterial morphological changes in PAS, as measured from top agar assays, has been emphasized in several studies since its re-discovery (7, 12). Remarkably, synergistic interactions were observed in the bacterial lawn regions immediately adjacent to the antibiotic inhibitory halos, after which plaque size returned to normal. The existence of an appropriate range of antibiotic concentrations where plaque enlargement takes place suggests the role of major physiological effects in the host that take place only at high but still sublethal drug concentrations. In the present study, the choice of liminal subinhibitory concentrations allowed us to reproduce such an effect in a controlled way. Comeau et al. studied synergy between filamentation-inducing antibiotics, and a phage (φMFP) infecting a uropathogenic *E. coli* strain (7). They suggested that the mechanistical basis for plaque enlargement was the net increase in phage progeny experienced upon antibiotic-induced filamentation. As a result, they proposed that an enlarged bacterial cytoplasm could provide more biosynthetic material and allowed the production of larger phage progenies. In a more recent work, an alternative explanation was suggested, based on the observation that filamentation delays the lysis of phage T4, with a consequent increase in burst size (12). Although both studies are based on taxonomically distantly related phages, they both agree on the correlation between cell filamentation and PAS on the one hand and on the increase in the burst size to explain lysis plaque enlargement. Through careful investigation of T5 and T7 viral cycles with dCas9-induced elongated cells (Fig. 4) we concluded that the impact of filamentation on phage replication ultimately leads to an increased burst size with both phages. Nevertheless, we observed phage-specific differences. In the case of phage T7, a clear increase of the burst size in the presence of dCas9/*ftsZ*-sgRNA-mediated filamentation was observed, without changing the latent period. As mentioned before, this behavior was observed with phage φMFP infecting *E. coli* in the presence of cefotaxime (7), and more recently with coliphage HK620 in the presence of cephalexin and ciprofloxacin (13). Conversely, upon bacterial filamentation, T5 displayed an increased burst size as a consequence of a delay in the latent period, which in turns aligns with the “delayed lysis” hypothesis proposed by Kim et al. According to this hypothesis, the concentration threshold required for the holin aggregation and disruption of the bacterial envelope increases as a consequence of membrane enlargement upon filamentation (12). T4 holin T belongs to the type III class of holins consisting in a single transmembrane domain with two soluble domains that protrude into the cytoplasm and periplasm and play a role in lysis-inhibition (24). Curiously enough, class III holins are only found in T4-like and T5-like phage families (25), which could explain the similar lysis-delay response upon bacterial filamentation.

In contrast, the drastic change in burst size observed in the presence of filamentation was not observed upon cell rounding (Fig. 4). In fact, no significant changes in the lytic cycle of phages T5 and T7 were detected under cell rounding mediated by dCas9/*mreB*-sgRNA expression. Since the timing and productivity of an individual viral cycle is not affected during the epidemic spread whether the cell retains its normal shape or become rounded, it is highly likely that the synergistic behavior arose from a change in the dynamics of virion diffusion from one host to the next one.

We have evidenced here that in the presence of cell rounding, whether it is caused by the addition of the antibiotic mecillinam or the repression of *mreB*, the microcolonies formed by *E. coli* tend to occupy less space due to a more compact spatial arrangement (Fig. 4). As a consequence, this abnormal aggregation pattern significantly reduces the fraction of space occupied by the bacterial populations and increases the free, unoccupied space between microcolonies in the matrix.

Ultimately, an increase in the free space between host clusters during phage spread can allow virions to freely diffuse farther away from the epidemic center between each cycle, allowing to travel longer distances before the encounter of a new non-infected bacterium. These results are in accordance with previous theoretical and experimental work suggesting that a reduction in phage-bacteria interaction could increase phage propagation speed (6, 26, 27). Thus, by reducing the likelihood of a phage encounter with a cluster of hosts, viral diffusion is the main parameter accounting for the increase in the propagation speed.

In the literature, lysis plaque expansion has been studied as a diffusion-reaction wave of phages over a bacterial lawn (27, 28). In early literature, common heuristic approaches for modelling plaque expansion rates were rather phenomenological (29). Later, Yin & McCaskill proposed a reaction-diffusion model consisting in diffusing phages that bind upon collision with susceptible, immobile and non-dividing bacteria (27). Upon irreversible binding, bacteria were infected and produced free virions. This model however, ignored bacterial division and the time spent by the phage replicating inside its host (the latent period). More recently, Fort and Mendez presented delayed reaction-diffusion models where the latent period is taken into account improving the predicted speed of plaque development (30). Such models are generally solved either analytically or numerically to obtain the speed of propagation of the traveling fronts as a function of burst size, latent time, intrinsic diffusion of phage and adsorption rate. We propose a new model including parameters we measured during this study like cell morphology as well as the physiological state of the bacteria as T7 allows lysis plaque expansion monitoring during both exponential and stationary phase. An interesting remark derived from our model is that the morphological parameters influencing phage propagation speeds are fundamentally cell length (*l*) and surface area-to-volume ratio (σ). The latter can be approximated to σ = s/v ≈ 1/w (the inverse of the cell width), in case of a spheric-cylindrical object. Ojkic et al. presented a model for the homeostasis of the bacterial cellular shape that explicits these two parameters, width and length as a function of the rate of accumulation of the division protein FtsZ ring and cell wall production (through the activity of the MreB protein) (31). They show that the width 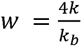 and length 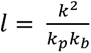, where k_p_ is the constant of accumulation of the division protein FtsZ, *k_b_* is the rate of cell wall production linked to MreB, and *k* is the rate of cell elongation. Accordingly, both ciprofloxacin and mecillinam might impact the dynamics of this process. Ciprofloxacin triggers the SOS response which impairs FtsZ assembly due to the production of the SulA inhibitor (32), thus decreasing the *k_p_*, and increasing cell length. On the other hand, mecillinam inhibits the transpeptidase site of its target PBP2, disrupting the normal functioning of MreB and its related proteins in the Rod system (33), this has a negative impact in *k_b_*, and consequently produce an increase in cell width. These studies provide a direct link between the morphological changes observed in the presence of both drugs.

The underlying mechanisms of phage-antibiotic synergy, defined as the increase in the size of a lysis plaque in a lawn of bacteria, started to be described in previous publications. However, the models presented thus far relied in the increased in phage fecundity as the main explanation for the synergistic contribution of antibiotics, ignoring additional contributions resulting from abnormal host morphologies. In this study, we demonstrate that dissimilar antibiotics seem to increase the size of a lysis plaque mainly by altering the shape of the bacteria that compose the lawn. In the model we propose, plaque enlargement is a consequence of imbalanced phage and bacteria interactions during plaque formation that consequently privileges phage spread to different degrees.

This study shows that plaque enlargement observed under phage-antibiotic synergy conditions is largely determined by antibiotic-induced changes in bacterial morphology, rather than by the specific molecular actions of antibiotics. By modifying the shape of the bacteria, for example by inducing filamentation or rounding, we observed a significant enlargement of the phage lysis plates, highlighting the role of bacterial morphology in improving phage propagation in structured environments. This finding moves us away from traditional models that focus on increased phage reproduction and instead highlights the critical impact of bacterial shape on phage epidemic propagation.

## Materials and Methods

### Bacterial strains, phages and culture conditions

Bacteriophage T5 was kindly provided by P. Boulanger’s team and bacteriophage T7 was obtained from the DSMZ collection. Bacterial host *E. coli* MG1655 used in this study originated from P. Genevaux’s collection. *E. coli* LC-E75 strain carrying a chromosomal copy of *dcas9* gene under the *P*_tet_ promoter was provided by David Bikard (Addgene #115925). *dcas9* encodes a *Streptoccocus pyogenes* Cas9 mutant inactivated in its endonuclease function (34). Liquid cultures were carried out at 37°C, 180 rpm in Lysogeny Broth (Thermo Fisher). For solid and semisolid media, 1.5 % or 0.5% agar (Sigma-Aldrich) were added, respectively. Antibiotics mecillinam (Sigma-Aldrich), ciprofloxacin (Merk), ceftazidime (Sigma-Aldrich), kanamycin (Sigma-Aldrich) 50 µg/mL and chloramphenicol (Sigma-Aldrich) 25 µg/mL were added at the indicated concentrations.

Anhydrotetracycline (ATc Sigma-Aldrich) was used at the indicated concentrations to induce *dCas9* gene expression.

### Plasmids and sgRNA spacers

Plasmid psgRNA coding for a single-guide RNA (sgRNA) used by dCas9 was a gift from David Bikard (Addgene plasmid # 11400). *mreB-* an *ftsZ-* targeting sgRNA were designed by hybridizing complementary primers carrying the following spacers: psgRNA-*ftsZ* 5’-CCTGAGGCCGTAATCATCGT-3’ and psgRNA-*mreB* 5’-GATATCAACCACCATAGAAC-3’. These spacers were flanked by two BsaI restriction sites and inserted into the psgRNA through Golden Gate assembly (NEB), thus generating *ftsZ*-sgRNA and *mreB*-sgRNA. The non-targeting sgRNA (sgRNA) refers to the unmodified plasmid. All three plasmids were transformed into chimiocompetent *E. coli* LC-E75 cells.

### Determination of antibiotics and inducer concentrations

In order to determine for each antibiotic the maximum concentration on plates that does not perturbs bacterial growth, we plated a well-mixed, log-phase bacterial culture in a 90 mm petri dish as follows: a volume of 5 mL of soft-agar (0.5 %) containing the bacterial culture at an initial optical density measured at 600 nm (OD_600_) of 0.1 was poured on top of a 20 mL layer of hard-agar (1.5 %) devoid from bacterium. Antibiotics were diluted in the hard-agar layer and the indicated concentrations were given for the total solid medium volume (5 mL soft-agar + 20 mL hard-agar).

Plates were then incubated overnight at 37°C. The bacterial lawn aspect was used to determine the maximum antibiotic concentration that neither significantly impact the opacity of the bacterial lawn nor produce grainy or heterogenous textures on the Petri dish surface. These sub-inhibitory concentrations were set at 15 ng/mL for ciprofloxacin, 120 ng/mL for ceftazidime, 200 ng/mL for mecillinam, 3 µg/mL for chloramphenicol and 2 µg/mL for kanamycin. *E. coli* cell rounding or filamentation was reproduced without antibiotics by repressing *mreB* and *ftsZ* gene expression, respectively. ATc was used to induce *dCas9* gene expression and the maximum concentration set at the value leading to morphological changes without eliciting significant cell death.

### Lysis plaque radii measurements

Phage lysis plaque expansion was monitored on plates by recording images with a stereomicroscope equipped with a 1X objective (Nikon SMZ800N). For single timepoint analysis, phage lysis plaques were left to propagate for 24 hours at 37°C then a single image was taken. For T7 plaque formation kinetics, we followed three distinct lysis plaques on a plate incubated at 37°C with image acquisition every 15 minutes. Plaques radii were determined using ImageJ plugin “Radial Profile Angle” (Paul Baggethun, Pittsburgh, PA) and selecting a radius of 1000 pixels (8.62 mm) centered on each lysis plaque. The resulting averaged intensity profiles were normalized using the formula 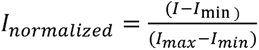. After normalization, a sigmoid curve was fitted to each measured plaque intensity profile using the non-linear least squares function from the R software and the following formula: 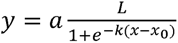 (where a and L are the lower and upper asymptotes, respectively, κ the growth rate or “steepness” of the curve, and *x*_0_ as the *x* value of the function midpoint). The radius value for a given plaque was considered to be the inflection point of the fitted sigmoid curve.

### Assessing antibiotics effect on bacterial morphology

A log-phase *E. coli* MG1655 culture was diluted to a final OD_600_ of 0.025 in 10 mL of LB medium, with or without addition of subinhibitory concentrations of antibiotics. After 2 hours incubation at 37°C under agitation at 180 rpm, a culture sample (OD_600_ ≈ 0.8) was fixed by diluting 1:1 in PBS buffer PFA 4% solution. Bacterial cells were imaged on an inverted phase-contrast microscope (Nikon TiE) using an oil immersion 100X NA 1.45 objective and Nikon’s NIS-Element software.

Large fields were captured to ensure statistical meaning of the analyzed population.

### Image analysis and morphological changes determination

Cell image analysis was performed using MicrobeJ (35) after treatment with Omnipose, a bacterial segmentation tool (36). From the output mask we then measured total length (*l*) from pole to pole and width (*w*) for each individual cell. To estimate bacterial surface area (*S*) and volume (*V*), each cell was modelled as a cylinder (length *l* – *w* and width *w*) capped with two hemispheres (diameter *w*) on each extremity:

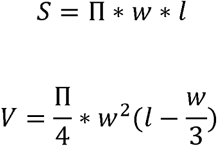

Statistical analysis was conducted in R (37) and figures were produced using the ggplot2 package (38).

### T5 and T7 latent period and burst size determination

Latent period (time between genome injection and release of the first virion by cell lysis) and burst size (number of virions produced per infected cell) were determined in liquid medium through the One-Step Growth Curve experiment (39). Log-phase cultures of *E. coli* LC-E75, carrying sgRNAs targeting either *mreB*, *ftsZ* or nothing, were infected at low Multiplicity Of Infection (MOI ∼0.001) at 37 °C and 180 rpm. In all cases, induction of dCas9 took place 2 hours prior to phage infection in order to allow the dCas9/sgRNA-mediated morphological change to take place. Five min post-infection, 100 µL culture samples were then regularly assayed (every 5.0 and 2.5 min for T5 and T7, respectively) and plated using the top-agar overlay method with *E. coli* MG1655 as indicator strain. The assessment of morphological changes induced by dCas9 expression was confirmed by microscopy on culture aliquots sampled shortly before infection. After overnight incubation of the plated samples, the number of Plaques Forming Units (PFU) at each timepoint was calculated and plotted against time.

### Experimental data fitting

In order to determine T7 lysis plaque expansion terminal velocities, experimental (*r - r*_0_) kinetics for each condition displayed in Figure S1 were fitted with a linear regression *y* = *a*_1_.*x* + *a*_0_ in the 15 – 20 h time range (terminal velocity c is equal to a_1_). Both a_O_ and a_1_ coefficients were computed according to the weighted least squares regression method. Since for each condition tested (r - r_O_) curve is the mean of three kinetics recorded on three individual lysis plaque, the standard deviation of the experimental data was used to calculate the terminal velocity c standard deviation.

The experimental squared relative terminal velocities with respect to cell width *w* (Fig. 5a) or cell length I (Fig. 5b) was fitted with a linear regression *y* = *a*_1_.*x* + *a*_0_ (Fig. 5a) or a 2^nd^ degree polynomial function *y* = *a*_2_.*x*^2^ + *a*_1_.*x* (Fig. 5b) in accordance with our theoretical model also using the weighted least squares regression method. The standard deviation of each a_i_ parameter was computed using the terminal velocities and morphological parameters standard deviations.

## Supporting information

Figure S1, Figure S2, Model description

## Acknowledgments

We are thankful to all present and past members of the Phages@LCB group for stimulating discussions and constant interest. We are grateful to Sylvain Gandon (CEFE) for challenging hypothesis and to Leon Espinosa (LCB) for advice and guidance in image analysis. We thank Cécile Giordano and the support staff in LCB for their constant and cheerful help. The Centre National pour la Recherche Scientifique (CNRS), Aix-Marseille Université (AMU) and the Institut de Microbiologie de la Méditerranée (IMM) support our research and provide exceptional on-campus facilities. The Mission for Transversal and Interdisciplinary Initiatives (MITI) from CNRS supported this work through the allowance of a doctoral fellowship to M.A.’s group as well as a collaborative grant “Adaptation of the living to its environment” in collaboration with Sylvain Gandon.

