## Supplementary material for "Antibiotic-Induced Morphological Changes Enhance Phage Propagation: A Mathematical Model of Plaque Formation in Structured Environments": Figure S1, Figure S2, Model description

**Antibiotic-induced morphological changes boost phage propagation through diverse mechanisms**

**This PDF file includes:**

Modeling of T7 lysis plaque propagation, including detailed equations corresponding to the mathematical model

SI References

Figures S1 to S2

**Other supporting materials for this manuscript include the following:**

Datasets S1

**Modeling of T7 lysis plaque propagation**

*Parameters*

*F*	Free phages fraction

*A*	Adsorbed phages fraction

*P*	Total phage concentration

*d*	Phages intrinsic diffusion coefficient (µm^2^.min^-1^)

*k*	Phage inactivation rate (min^-1^)

*µ*	Biomass growth rate (h^-1^)

*V*	Biomass volume (µm^3^)

$l$	Cell length (µm)

$w$	Cell width (µm)

$s$	Cell surface area (µm^2^)

$v$	Cell volume (µm^3^)

*s*	surface area-to-volume ratio (µm^-1^)

*a*	Phage adsorption rate (cell^-1^.µm^-2^.min^-1^)

*g*	Phage replication rate per unit cell length (min^-1^.µm^-1^)

*b*	Phage release rate (min^-1^)

*Equations*

In our system, the total phage concentration $P$ dynamics is a function of the dynamics of two “species” of phages within the sample, namely, “free” phages (denoted by $F$) and “absorbed” phages (denoted by *A*). “Free” phages diffuse freely through the substrate with an intrinsic diffusion coefficient of $\delta$ ($\mu m^{2}min^{-1}$) and become inactive at a rate $\kappa\left( min^{-1} \right)$. In the presence of *E. coli* susceptible cells, free phages adsorb onto the cell surface with a rate proportional to the cell surface area (*s*) (10). We consider the biomass volume *V* and the cell volume $v$to write the rate of adsorption as $\alpha\frac{s}{v}V$, with an adsorption rate per surface area per cell $\alpha$ ($cell^{-1}\mu m^{-2}min^{-1})$. We assumed adsorbed phages replicate within the cell with a rate proportional to the cell length ($l$) and rate per unit length $\gamma$ ($min^{-1}\mu m^{-1}$) (1). Phages are released from the cell at a constant rate $\beta$ ($min^{-1}$). Then, the dynamical equations for the free and adsorbed phages are written as:

$F_{t}=\delta F_{xx}-\alpha\frac{s}{v}VF+\beta A-\kappa F$	(Eq. 1)

$A_{t}=\alpha\frac{s}{v}VF+\gamma lA-\beta A$	(Eq. 2)

$P_{t}=F_{t}+ A_{t}$	(Eq. 3)

where the subscripts *t* and *xx* represent the derivative with respect to time, and second derivative in space, respectively. Under the assumption that the diffusion is slower than the adsorption and release of the phages, $A_{t}\to0$. Therefore, we may simplify Eq. 1 and Eq. 2 as:

$A=\frac{\lambda\sigma V}{1+\lambda\sigma V}P$	(Eq. 4)

$F=\frac{1}{1+\lambda\sigma V}P$	(Eq. 5)

where $\lambda=\frac{\alpha/\beta}{1-\gamma l/\beta}$. Adding the above two equations and writing the dynamical equation for the total phage and considering that the biomass volume $V$ grows slowly compared to the above dynamic equations, we simplify the total phage concentration $P$ as:

$P_{t}=\frac{\delta}{1+\sigma\lambda V}P_{xx}+\gamma l\frac{\sigma\lambda V}{1+\sigma\lambda V}P-\frac{\kappa}{1+\lambda\sigma V}P$	(Eq. 6)

where $\sigma=s/v$. Then, we look for a travelling wave front solution of the form $P\left( z \right)=P\left( x-ct \right)$. We thus obtain:

$-c\left[ P\left( z \right) \right]_{z}=\frac{\delta}{1+\sigma\lambda V}P\left( z \right)_{zz}+\gamma l\frac{\sigma\lambda V}{1+\sigma\lambda V}P\left( z \right)-\frac{\kappa}{1+\lambda\sigma V}P\left( z \right)$	(Eq. 7)

$P\left( z \right)_{z}=Q\left( z \right)$(Eq. 8a)

$Q\left( z \right)_{z}=-\frac{c}{\delta}\left( 1+\sigma\lambda V \right)Q\left( z \right)-\gamma l\sigma\lambda VP\left( z \right)/\delta+\kappa P\left( z \right)/\delta$	(Eq. 8b)

The Jacobian *J* near steady state is then:

$J=\left[ \left[ 0,\frac{\kappa}{\delta}-\frac{\gamma\lambda l\sigma V}{\delta} \right] \right],\left[ 1,-\frac{c\left( \lambda\sigma V+1 \right)}{\delta} \right]$	(Eq. 9)

with eigenvalues *eig*:

$eig=-\frac{c\left( \lambda\sigma V+1 \right)\pm\sqrt{\left( c\lambda\sigma V+c \right)^{2}+4\delta\left( \kappa-\gamma\lambda l\sigma V \right)}}{2\delta}$	(Eq. 10)

Real roots exist under the condition that:

$c^{2}\geq4\delta\frac{\gamma l\lambda\sigma V-\kappa}{\left( 1+\lambda\sigma V \right)^{2}}$	(Eq. 11)

leading to the expression for the minimum speed of the travelling wavefront solution.

In a model where a bacterium is represented by a cylinder caped by two hemispheres, cell surface area $s$, cell volume $v$ and the surface area-to-volume ratio s are:

$s = \pi w \left( l - w \right)+ 4 \pi\left( \frac{w}{2} \right)^{2}$	(Eq. 12)

$v = \pi\left( \frac{w}{2} \right)^{2} \left( l - w \right)+ \frac{4}{3} \pi$	(Eq. 13)

$\left( \frac{w}{2} \right)^{3}\sigma=\frac{s}{v}= 12 \frac{l}{w \left( 3l - w \right)}\sim\frac{const.}{w}$	(Eq. 14)

Since we modelled a bacterium as a cylinder caped by two hemispheres, the surface area-to-volume ratio is inverse of the width of bacteria, *i.e.*, $\sigma\sim1/w$. The stability analysis of this solution to travelling wavefront solution of the form $P\left( x-ct \right)$ yields the minimum speed of the wavefront $\left| c \right|$ in the form of:

$\left| c \right|\geq2\sqrt{\delta\beta}\frac{\sqrt{\frac{\gamma l}{\beta}\lambda\sigma V-\frac{\kappa}{\beta}}}{1+\lambda\sigma V}$	(Eq. 15)

The solution thus exists under the conditions that replication rate is slow, $\gamma l<\beta$ and the virion inactivation rate is small, $\kappa<\gamma l\lambda\sigma V$. The speed is shown to be a function of a characteristic speed $\sqrt{\delta\beta}$, morphological parameters $\frac{\gamma l}{\beta},\sigma$, phage-related parameter $\alpha/\beta$ and the biomass volume $V$ encountered by the travelling wave. We thus defined from Eq. 4 the dimensionless speed $c/\sqrt{\delta\beta}$ as:

$c/\sqrt{\delta\beta}=2\frac{\sqrt{\frac{\gamma l}{\beta}\lambda\sigma V-\frac{\kappa}{\beta}}}{1+\lambda\sigma V}$	(Eq. 16)

By modelling the biomass in-front of the traveling wave as a logistic growth with growth rate $\mu$ we can obtain a time-dependent speed by defining $V\left( t \right)$:

$V\left( t \right)=\frac{V_{max}}{1+\left( \frac{V_{max}}{V_{0}}-1 \right)e^{-\mu t}}$	(Eq. 17)

where $V_{max}$ is the carrying capacity and $V_{0}$ is the initial biomass volume.

In the case where phage deactivation is negligible ($\kappa\ll\gamma l$), which seems to be the case for T7 phage, the speed of the travelling wavefront $\left| c \right|$ (Eq. 15) can be simplified as:

$\left| c \right|\geq2\frac{\sqrt{\delta\beta} \sqrt{\frac{\gamma l}{\beta}\lambda\sigma V}}{1+\lambda\sigma V}$	(Eq. 18)

Furthermore, when the host reaches biomass capacity $V_{max}$ (and assuming $\lambda\sigma V_{max}\gg1$), the travelling wavefront reaches a terminal velocity $c$:

$c\sim2\sqrt{\frac{\delta\gamma l}{\lambda\sigma V_{max}}}\sim2\sqrt{\frac{\delta\gamma}{V_{max}}} \sqrt{\frac{lw}{\lambda}}$	(Eq. 19)

Thus, the square of speed in each experimental case $c_{T}$ (*T* for antibiotic-treated cells) relative to the speed of untreated host $c_{u}$ (*u* for untreated cells) is:

$\left( \frac{c_{T}}{c_{u}} \right)^{2}=\frac{l_{T}w_{T}}{l_{u}w_{u}}\frac{1-\frac{\gamma l_{T}}{\beta}}{1-\frac{\gamma l_{u}}{\beta}}$	(Eq. 20)

When cell length $l$ is constant (*e.g.*, with mecillinam), Eq. 20 can be simplified as a first-degree polynomial equation with a null intercept:

$\left( \frac{c_{T}}{c_{u}} \right)^{2}=\frac{w_{T}}{w_{u}}=a_{1}*w_{T}$	(Eq. 21)

with $a_{1}= \frac{1}{w_{u}}$.

When cell width $w$ is constant (*e.g.*, with ciprofloxacin), Eq. 18 can be simplified as a second-degree polynomial equation with a null intercept:

$\left( \frac{c_{T}}{c_{u}} \right)^{2}=\frac{1}{l_{u}(1-\frac{\gamma}{\beta}l_{u})}l_{T}*\left( 1-\frac{\gamma}{\beta}l_{T} \right)=a_{1}*l_{T}*(1-a_{2}*l_{T})$	(Eq. 22)

with $a_{1}= \frac{1}{l_{u}(1-\frac{\gamma}{\beta}l_{u})}$ and $a_{2}= \frac{\gamma}{\beta}$.

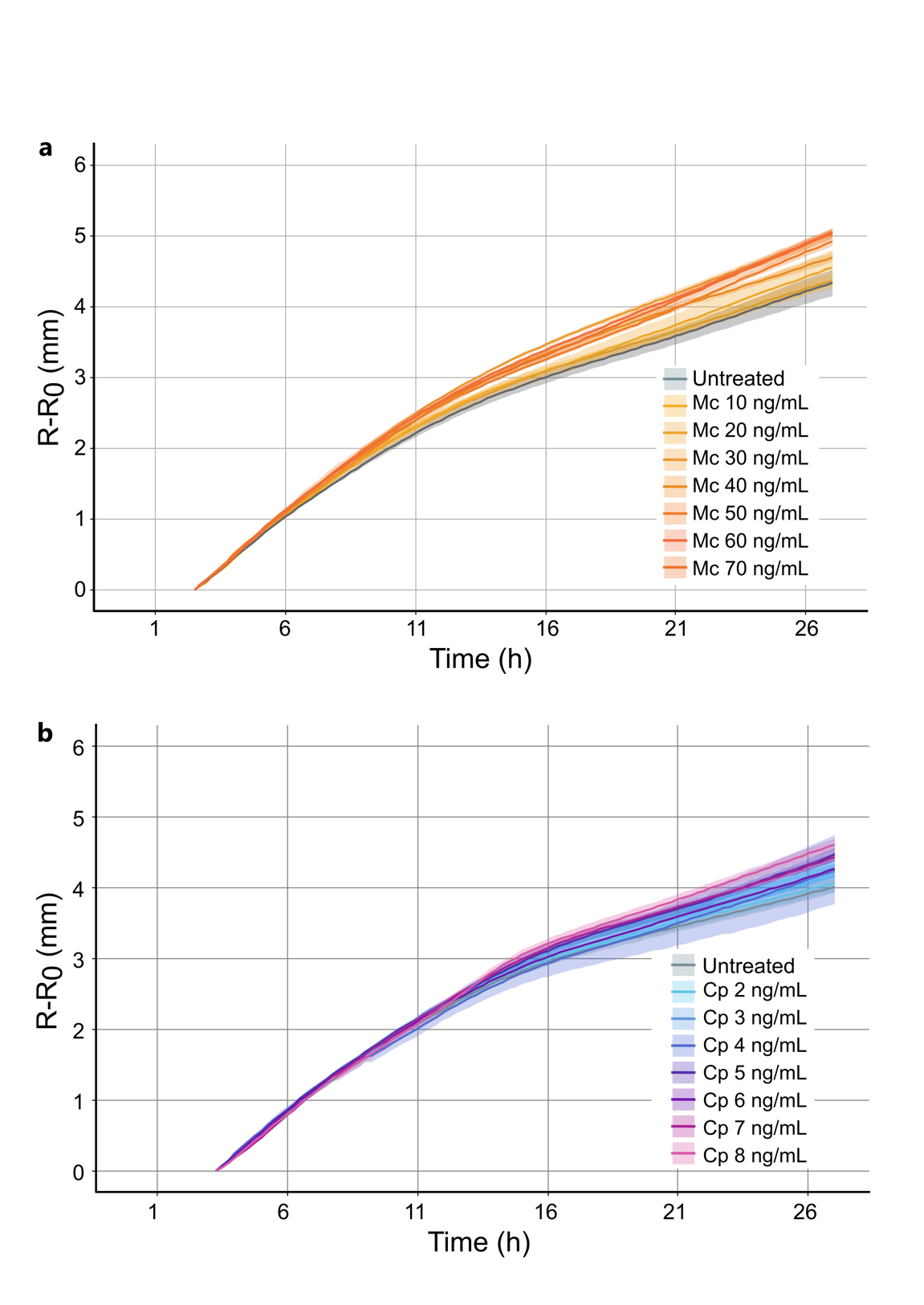

**Figure S1: Propagation profiles of phage T7 epidemics under PAS conditions.** Radius increase of T7 lysis plaques between 3 and 28 hours post infection in the presence of increasing concentration of mecillinam (a) or ciprofloxacin (b). N= at least 3 independent plaques were recorded per condition. The shadowed areas represent the standard error of the mean of each curve (S.E.M).

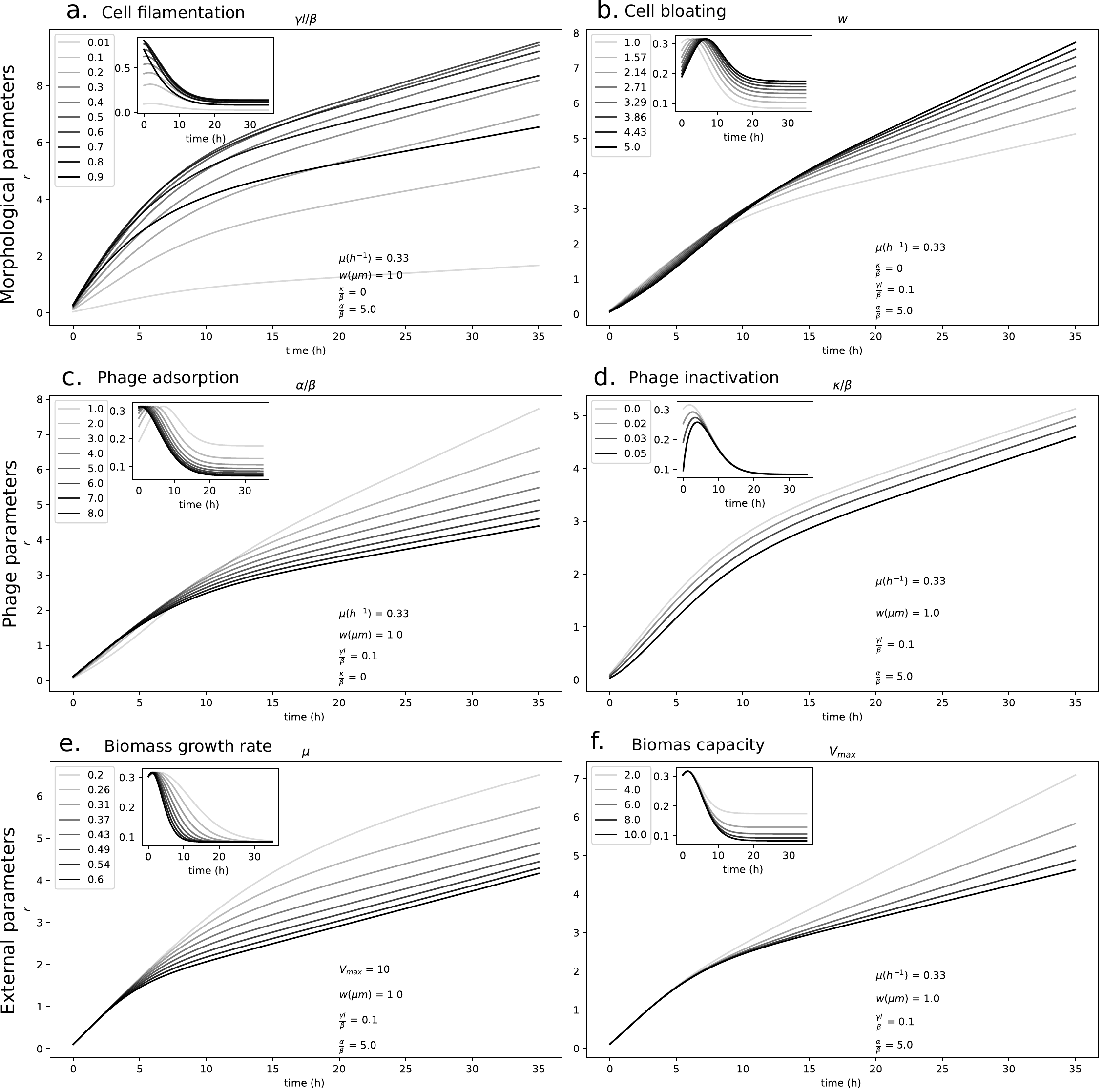

**Figure S2: Effect of model parameters on phage lysis plaque expansion.** Phage lysis plaque radius $r$ expansion kinetics (main panels) are obtained by integration of velocity changes over time (insets). The velocity scaling factor $\sqrt{\delta\beta}$ is kept constant at value 1.0 (Eq. 18). **(a, b)** *Effect of cell morphological parameters.* Cell length $l$ (**a**) and cell width $w$ (**b**). **(c, d)** *Effect of phage parameters.* Phage adsorption (**c**) and phage inactivation (**d**). **(e, f)** *Effect of external parameters.* Biomass growth rate $\mu$ (**e**) and biomass capacity $V_{max}$ (**f**).
